# Within and among-population variation in vital rates and population dynamics in a variable environment

**DOI:** 10.1101/028662

**Authors:** Simone Vincenzi, Marc Mangel, Dusan Jesensek, John Carlos Garza, Alain J Crivelli

## Abstract

Understanding the causes of within-and among-population differences in vital rates, life histories, and population dynamics is a central topic in ecology. To understand how within-and among-population variation emerges, we need long-term studies that include episodic events and contrasting environmental conditions, data to characterize individual and shared variation, and statistical models that can tease apart population-, shared-, and individual contribution to the observed variation.

We used long-term tag-recapture data and novel statistical and modeling techniques to investigate and estimate within-and among-population differences in vital rates, life histories and population dynamics of marble trout *Salmo marmoratus*, a endemic freshwater salmonid with a narrow range. Only ten populations of pure marble trout persist in headwaters of Alpine rivers in western Slovenia. Marble trout populations are also threatened by floods and landslides, which have caused the extinction of two populations in recent years. We estimated and determined causes of variation in growth, survival, and recruitment both within and among populations, and evaluated trade-offs between them. Specifically, we estimated the responses of these traits to variation in water temperature, density, sex, early life conditions, and the occurrence of extreme climatic events (e.g., flash floods and debris flows).

We found that the effects of population density on traits were mostly limited to the early stages of life and that individual growth trajectories were established early in life. We found no clear effects of water temperature on survival and recruitment. Population density varied over time, with flash floods and debris flows causing massive mortalities and threatening population persistence. Apart from flood events, variation in population density within streams was largely determined by variation in recruitment, with survival of older fish being relatively constant over time within populations, but substantially different among populations. Marble trout show a fast-to-slow continuum of life histories, with slow growth associated with higher survival at the population level, possibly determined by food conditions and age at maturity.

Our work provides unprecedented insight into the causes of variation in vital rates, life histories, and population dynamics in an endemic species that is teetering on the edge of extinction.

## 1 Introduction

Understanding the causes of within-and among-population variation in vital rates (e.g., survival, body growth), life histories, and population dynamics is a central topic in ecology (Frederiksen et al. 2014). This task is even more crucial when the species investigated are of conservation concern, since a robust understanding of the effects of environmental (e.g., weather, food) and population (e.g., density-dependent) processes on these traits and dynamics, and on risk of local population extinction, is critical for population forecasting and species management (Morris and Doak 2002).

Comparative quantitative studies of population-, group and individual-level traits covering a substantial part of a species’ geographic range are very rare (Frederiksen et al. 2005, Suryan et al. 2009). In order to understand how within-and among-population variation in vital rates and life histories of organisms emerge we need (1) long-term studies that include contrasting environmental conditions and rare, but influential, events (Elliott 1994), (2) data for the estimation and characterization of individual and shared (i.e., among groups) variation (Thomson et al. 2009), and (3) statistical models that can tease apart population, shared, and individual contributions to the observed variation in demography, life histories, and population dynamics (Letcher et al. 2015). In addition, given the ample opportunity for exploratory analyses when data from many potential explanatory variables are collected, it is crucial to organize the analyses of this variation around well-defined and theory-based hypotheses that limit the “researcher degrees of freedom” (Simmons et al. 2011).

Habitat variation plays a major role in determining intra-specific differences in life histories, behavior and physiology of organisms (Suryan et al. 2009, Jonsson and Jonsson 2011). Habitat factors can be either extrinsic (e.g., weather, predators, food availability) or intrinsic (e.g., population density). As habitats differ geographically and temporally, so can the distribution of vital rates, life histories, phenotypic characters, and genetic structure of conspecific populations (Linhart and Grant 1996). Long-term studies allow us to quantify the relationships between demography and habitat, both because of increased statistical power and because they are more likely to include periods with contrasting environmental conditions (Clutton-Brock and Sheldon 2010). Long-term data are also necessary to understand ecological processes that are driven by episodic, but influential, events (Smith 2011).

One such example of episodic events are extreme climatic events (Smith 2011), which are, by definition, rare and influential. Their ecological and genetic effects include dramatic demographic crashes or extinction of populations or species (Piessens et al. 2009), genetic bottlenecks (Shama et al. 2011), changes in age-and size-structure (Chan et al. 2005), and shifts in the phenology of plant and animal species (Jentsch et al. 2009). Since climate change is predicted to increase the frequency and intensity of extreme climatic events (IPCC 2007, 2012), population responses to such events need to be carefully investigated for conservation purposes (Vincenzi 2014; Ohlberger and Langangen 2015).

Within populations, organisms often differ in the ability to acquire resources, in their life-history strategies, and in their contributions to the next generation (Lomnicki 1988). This variation may result from complex interactions between genetic, environmental and population factors, and can have substantial consequences for both ecological and evolutionary dynamics (Lomnicki 1988, Pelletier et al. 2007, Coulson et al. 2010). Longitudinal data, such as those provided by tag-recapture studies, have greatly facilitated the estimation of individual and group (i.e., sex, year-of-birth cohort) variation in reproductive success, survival, and growth (Thomson et al. 2009).

Individual and group variation also affects the estimation of parameters in population and life-history models, which may translate to incorrect predictions and inference (Pfister and Stevens 2003, Coulson et al. 2009, Smallegange and Coulson 2013). When multiple measurements from the same individuals are used for the estimation of model parameters, random-effect models (Gelman and Hill 2006) provide an intuitive framework for estimating or taking into account individual heterogeneity.

In this work, we used long-term tag-recapture data and powerful statistical and modeling techniques (Laake et al. 2013, Vincenzi et al. 2014b) to estimate and determine the causes of within-and among-population differences in vital rates, life histories, and population dynamics of marble trout *Salmo marmoratus* (Cuvier), a freshwater salmonid species endemic to rivers tributary to the upper Adriatic Sea.

Marble trout is a species of great conservation concern, given its restricted geographical distribution and the risk of hybridization with alien brown trout *Salmo trutta* L. Presently, the only (eight) genetically pure natural populations of marble trout are located in the Adriatic basin of Western Slovenia (Fig. 1) (Berrebi et al. 2000, Crivelli et al. 2000, Fumagalli et al. 2002), persisting above barriers that have prevented the upstream movement of brown trout or marble-brown trout hybrids living in the lower stream reaches (Sušnik et al. 2015). These populations are highly genetically differentiated, with intraspecific pairwise *F*_ST_ values among the highest ever reported (Fumagalli et al. 2002). To increase the number of viable marble trout populations, two other populations were created in previously fishless streams by translocation of the progeny of wild trout from the pure populations. There is little potential for spontaneous colonization of new habitats through dispersal or re-colonization after local extinctions. Marble trout populations are also threatened by extreme climatic events, such as flash floods, debris flows, and landslides that further increase the risk of species extinction. A conservation program for marble trout was started in Western Slovenia in 1993 (Crivelli et al. 2000); since then, the ten populations have been the subject of an intensive monitoring and tagging program. Since the start of this conservation-oriented program, extreme climatic events have caused the extinction of two populations, as well as population crashes in multiple other populations (Vincenzi et al. 2008d, 2010a, 2014a).

**Fig. 1.**
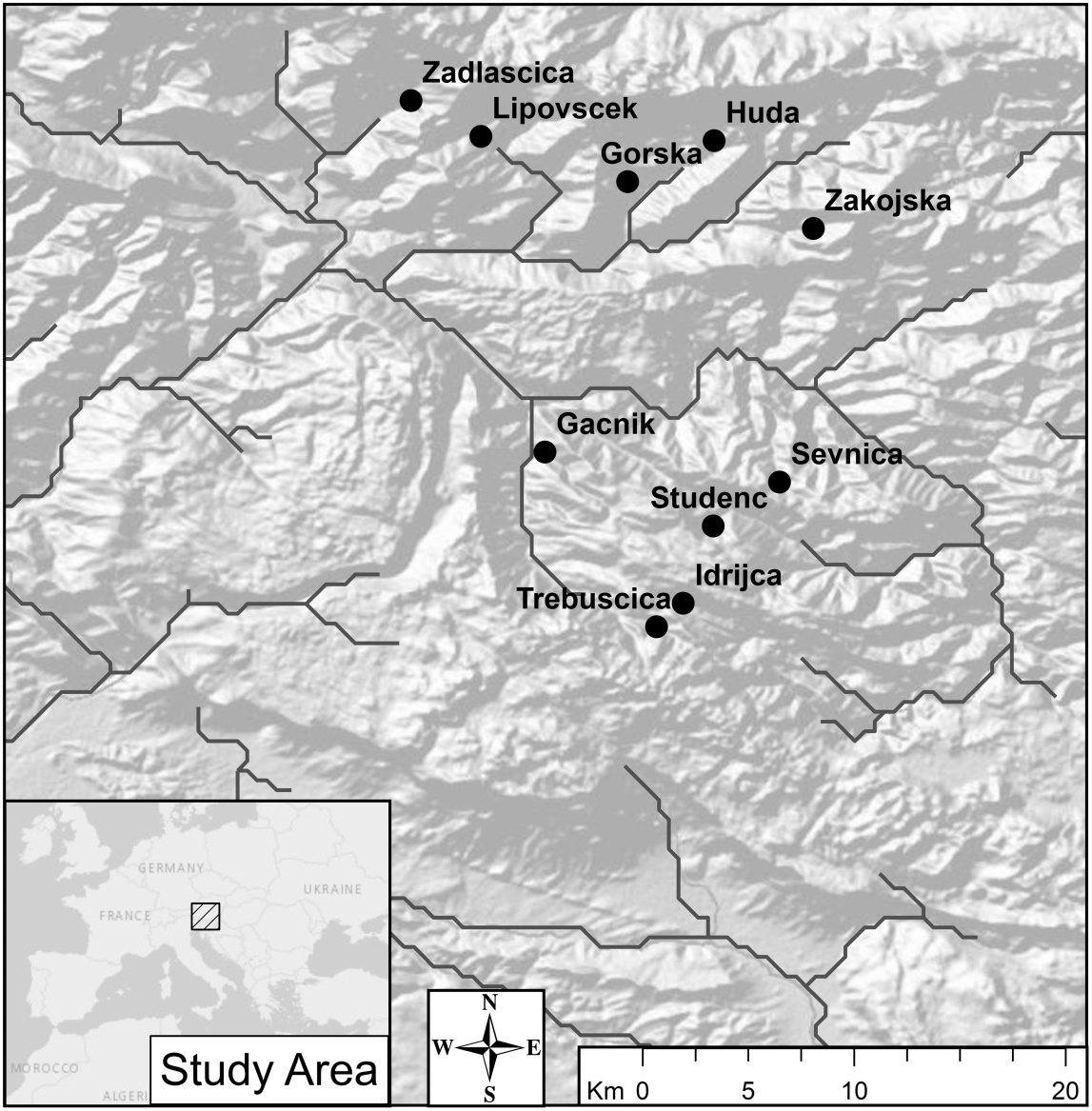
Remnant (Zadla, Lipo, Huda, Sve, Stu, Idri[LIdri, UIdri]) and newly created (Zak and Gac) populations of marble trout in Western Slovenia. Gorska was obliterated by a flash flood in 2004.

In this work, we estimated and determined causes of variation and trade-offs in growth, survival, and recruitment within and among marble trout populations. We use data from the intensive long-term tag recapture to investigate the responses of survival and growth rates to variation in water temperature, population density, sex, environmental conditions, and the occurrence of extreme climatic events. In particular, we tested hypotheses on determination of life histories early in life, the strength of cohort effects and their role in population dynamics, the relative role of stock size and environmental conditions in determining recruitment, life-history variation among populations and its determinants, and the mechanisms allowing population persistence after collapses in population size.

Our study includes all extant, pure marble trout population (Sušnik et al. 2015) and is thus one of the few studies that encompass a species’ entire geographic range (Valladares et al. 2014). Our work also provides a framework for the estimation of individual-, group-, and population-specific vital rates and life histories, using tag-recapture data and powerful analytical techniques for integrating individual-and/or group-level variation in growth and survival into the estimation of population-and species-level traits, as well evaluating the effects of extreme climatic events affecting them. In addition, the insights from this work provide clear guidance for conservation action that will improve the prospects of this species for avoiding extinction.

## 2 Material and Methods

### 2.1 Species and study area

There are eight, extant, natural, genetically pure marble trout populations, all isolated and separated from the downstream hybrid trout by impassable waterfalls. These populations live in headwater streams in the basins of the Soca, Baca, and Idrijca Rivers (the latter two are tributaries of the Soca/Isonzo River): Huda [Huda], Lower Idrijca [LIdri], Upper Idrijca [UIdri], Lipovesck [Lipo], Studenc [Stu], Svenica [Sve], Zadlascica [Zadla], Trebuscica [Trebu] (Fig. 1). All abbreviations and symbols used herein are found in Table S1. The populations of Lower and Upper Idrijca are separated by a dam that partially isolates them; in LIdri, marble trout live in sympatry with introduced rainbow trout, while in UIdri rainbow trout are absent (Vincenzi et al. 2011). All the other populations are in streams where marble trout is the only fish species and are located in pristine and remote locations without any regulated fishing, poaching, or presence of predators. Two additional populations of pure marble trout have been established in stretches of fishless streams - Zakojska [Zak] in 1996 and Gacnik [Gac] in 1998 - by translocation of age-1 fish (parental cohort) raised in the fish farm that were the progeny of fish from Zadla for Zak and from Trebu (females) and Lipo (males) for Gac (Crivelli et al. 2000). Fish first hatched in the streams in 1998 and 2000 in Zak and Gac, respectively.

Marble trout feed on benthic and terrestrial invertebrates, and on smaller marble trout (gape size is ~80%, Dusan Jesensek, *personal communication*) (Musseau *et al. in press*).

#### 2.1.1 Sampling

Populations were sampled either annually in June (Zak, Gac) or September (Zadla, Trebu, Sve) or bi-annually in both months (Huda, LIdri, UIdri, Stu, Lipo) (Table S2 and Fig. S1). Tagging started in different years for different populations: 1996 - Zak; 1998 - Gac; 2002 - Huda; 2004 - LIdri and UIdri; 2006 - Stu, Lipo, Trebu; and 2008 - Sve. Fish were captured by electrofishing and fork length (*L*) and weight recorded to the nearest mm and g, respectively. If captured fish had *L* > 110 mm, and had not been previously tagged or had lost a previously applied tag, they received a Carlin tag (Carlin 1955) and age was determined by reading scales. The adipose fin was also removed from all fish captured for the first time, including those not tagged due to small size. Fish are aged as 0+ in the first calendar year of life, 1+ in the second year and so on. Sub-yearlings are smaller than 110 mm in June and September, so fish were tagged when at least aged 1+. Males and females are morphologically indistinguishable in either June or September. The oldest fish were sampled in Gac (age 14) and LIdri (age 16), while in all other populations the oldest fish were age 9 or 10, with the exception of Stu, where the oldest sampled fish was age 6.

Unless otherwise noted, in the analyses shown below we used data from the start of tagging up to June (for Zak and Gac) or September (all other populations) of 2014.

#### 2.1.2 Environmental data

Stream length and topographical features were obtained from surveyor field investigations and from available GIS (Source: Surveying and Mapping Authority of the Republic of Slovenia) (Table S2). Annual rainfall in the area is between 2500 and 3000 mm. ONSET temperature loggers recorded mean daily water temperature in each stream. We used water temperature data to calculate growing degree-days (*GDDs*) with the formula *GDD* = *T*_mean_ - *T*_base_, where *T*_mean_ is the mean daily water temperature and *T*_base_ is the base temperature below which growth and development are assumed to stop. We set *T*_base_ at 5 °C as commonly used for salmonids (Chezik et al. 2014). Results are typically insensitive to variation in *T*_base_ over the 0-10 °C range (Chezik et al. 2014). *GDDs* are the sum of the days in which growth is possible during a specific time period (Chezik et al. 2014). Water flow rates data for the streams in which marble trout live have never been collected and thus water flow rates could not be used in any of the analyses presented below. All streams in which marble trout live are spring-fed.

#### 2.1.3 Floods

Streams of Western Slovenia are frequently affected by flash floods and debris flows (i.e. fast-moving landslides following intense rainfall) causing massive mortalities (Vincenzi et al. 2014a). Floods occurred in most of the streams in the fall of 2004, with noticeable effects on density of fish in Stu, Sve, and Lipo (before the start of tagging, see Table S2). After the start of tagging, severe fall floods occurred in Zadla (2007 and 2012), Zak (2007), Lipo (2007 and 2009), and Sve (2012) causing habitat modification, such as reshaping of the streambed, movement of boulders, and uprooting of trees. In addition, the floods of 2004 wiped out the population of Gorska, and the population of Predelica (Fumagalli et al. 2002) was extirpated by a landslide in 2000.

### 2.2 Density

We estimated density of 0+ fish in September for all populations except Zak and Gac, since fish emerged a few days or weeks before the June sampling of these populations. We estimated density of fish older than 0+ for three size classes: *L* ≤ 200 mm, *L* > 200 mm, or all fish older than 0+, using a two-pass removal protocol (or three-pass when needed), after confirming that a three-pass removal provided the same results as the two-pass removal.

We used total stream surface area for the estimation of fish density. The movement of marble trout is very limited, and most fish were sampled within the same 50-100 m stream reach throughout their entire lifetime (Vincenzi et al. 2008c). In addition, some of the populations are clearly circumscribed by impassable waterfalls both upstream and downstream. As such, it was possible to effectively survey the entire populations of Zak, Gac, Huda, while for the others only a fraction of the population was sampled. For the latter populations (excluding LIdri, in which only the tagging sector was used), fish were sampled and tagged in one sector, and data from another sector in which fish were sampled and not tagged were used to produce more accurate estimates of density. Huda is a very small population, with total number of sampled fish older than 0+ ranging from 23 to 118 since the start of tagging. The number of fish older than 0+ sampled in Zak ranged from 11 (in year 2008) to 500 (1996); in Gac from 212 (2000) to 1415 (2003) fish. In some populations, the estimation of density started before tagging (Table S2). We tested for recruitment-driven population dynamics by estimating correlations between density of 0+ fish (*D*_0+_) in September and density of fish older than 0+ (*D*_>0+_) one year later.

### 2.3 Growth and body size

In order to characterize size-at-age and growth trajectories, we modeled variation in size at first sampling (i.e., 0+ in September), lifetime growth trajectories, and daily growth between sampling occasions. We also estimated the repeatability of body size to test the hypothesis of maintenance of size hierarchies through marble trout lifetime.

#### 2.3.1 Variation in size at age 0+

We used an ANCOVA model to model the variation in mean length of cohorts at age 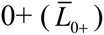 using *Stream*, *D*_>0+_, and *GDDs* (up to August included) and their interactions as candidate predictors. Following Vincenzi et al. (2008a, 2010b) and studies on density dependence of growth in salmonids (summarized in Jonsson and Jonsson 2011), we pooled together population-specific data and log-transformed both 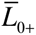 and *D*_>0+_. We excluded cohorts with < 3 fish sampled in September. We carried out model selection with the *MuMIn* package (Barton 2013) for R (R Development Core Team 2014), using the Akaike Information Criterion, AIC (Akaike 1974, Symonds and Moussalli 2010) as a measure of model fit. We considered that models had equal explanatory power when they differed by less than 2 AIC points (Burnham and Anderson 2002). We also tested whether there was density-dependence in mean condition factor of fish aged 0+ in September 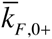 - where condition factor 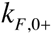 of individual *i* is 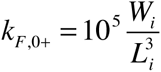 (*L_i_* in mm and *W_i_* = weight in g) (Froese 2006) - using the same ANCOVA model described for 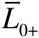.

#### 2.3.2 Lifetime growth trajectories

The standard von Bertalanffy model for growth (vBGF; von Bertalanffy 1957) is

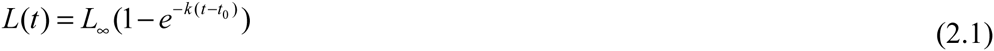

where L_∞_ is the asymptotic size, *k* is a coefficient of growth (in time^−1^), and *t*_0_ is the hypothetical age at which length is equal to 0.

We used a recently developed formulation of the vBGF specific for longitudinal data in which L_∞_ and *k* may be allowed to be a function of shared predictors and individual random effects (Vincenzi et al. 2014b). In the estimation procedure, we used a log-link function for *k* and *L*_∞_, since both parameters must be non-negative. We set

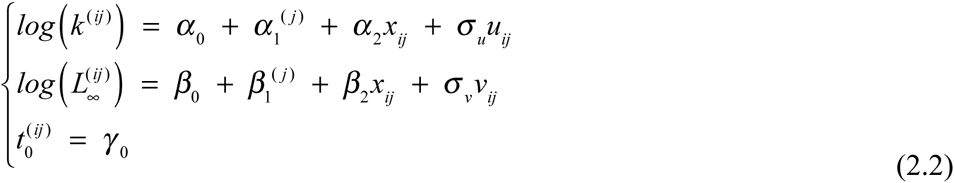

where *u* ~ *N* (0,1) and *v* ~ *N* (0,1) are the standardized individual random effects, *σ_u_* and *σ_v_* are the standard deviations of the statistical distributions of the random effects, and the other parameters are defined as in Eq. (2.1). The continuous predictor *x_ij_* (i.e., population density in the first year of life or temperature, as explained below) in Eq. 2.2 must be static (i.e., its value does not change throughout the lifetime of individuals).

We then assume that the observed length of individual *i* in group *j* at age *t* is

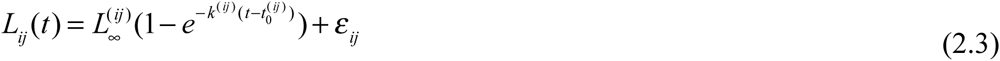

where *ε_ij_* is normally distributed with mean 0 and variance 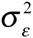.

Since the growth model operates on an annual time scale and more data on tagged fish were generally available for September, we used September data for the populations that were sampled twice a year. Following Vincenzi *et al.* (2014b), we included two potential predictors of *k* and *L*_∞_, (*i*) cohort (*Cohort*) as a group (i.e., categorical) variable (*α*_1_ and *β*_1_ in Eq. 2.2), and (*ii*) population density (of fish older than 0+) in the first year of life (*D*_>0+,_*_born_*) as a continuous variable (i.e., *x_ij_* Eq. 2.2) (Vincenzi *et al.* 2008b). In addition, we introduced (*iii*) sex (*Sex*) and (*iv*) *GDDs* in the first year of life as potential predictors of *k* and *L*_∞_. For the two newly created populations (Zak and Gac), previous work showed that the parental cohort born and raised in a hatchery grew faster and reached a bigger mean length-at-age than fish born in the stream (Vincenzi et al. 2008c). Thus, we also used (*v*) a binary predictor (*Coh*_p_) for *k* or *L*_∞_ to separately estimate parameters of the vBGF for the parental cohort and the cohorts born in the stream (pooled together). For the populations of Zadla and Lipo, we used (*vi*) a binary predictor (*Coh*_fl_) for *k* or *L*_∞_ to estimate separately vBGF’s parameters for cohorts born before the flood (pre-2007) and after the flood. For Zak, we used (*vii*) a categorical predictor (*Coh*_p.fl_) to differentiate the parental cohort, and cohorts born before and after the 2007 flood.

For each population, we also estimated the correlation between *k* and *L*_∞_ in models with no predictors for either *k* or *L*_∞_ to test the hypothesis that growth trajectories tend to not cross (strong positive correlation, i.e. size ranks tend to be maintained through the lifetime) or cross (strong negative correlation, i.e. size ranks tend not to be maintained through the lifetime) through a fish’s lifetime (Vincenzi et al. 2014b).

##### Datasets for the analysis of lifetime growth trajectories

When using shared *k* and *L*_∞_ among all fish in a population (i.e. no predictors for either *k* or *L*_∞_) or when using *Coh*, *Coh*_p_, *Coh*_fl_, and *Coh_p.fl_* as their predictors we used the whole datasets (datasets DataW) for every population.

We were able to assign sex using molecular techniques (Yano et al. 2013) for all fish in the populations of Trebu, Zadla, and Sve, and a subset of the fish in the population of Zak (1033 out of 1738 tagged individuals) and were thus able to use *Sex* as a predictor of *k* or *L*_∞_ in these populations (with dataset DataS for Zak).

There were fish in every population that were born before *D*_>0+,_*_born_* was measured and temperature data were available. For instance, in Huda the oldest-born tagged fish was born in 1997, while *D*_>0+,_*_born_* and water temperature were first estimated and recorded in 2002.

Since we wanted to compare the explanatory power of the growth models when *D*_>0+,_*_born_*, *GDDs*, *Coh, Coh*_p_, and *Coh*_fl_ were used as predictors of vBGF’s parameters, we used a subset of the whole datasets (datasets Data_D_) when using *D*_>0+,_*_born_* and *GDDs* as potential predictors. In each analysis, we used AIC to select the best model.

### 2.4 Repeatability of size

We estimated the repeatability of body size (Wilson et al. 2010), i.e. the proportion of total variance in body size that can be attributed to among-individual variation. High repeatability indicates that size ranks tend to be maintained throughout fish lifetime. To avoid the potential inflation of repeatability due to thinning of populations at older ages, we estimated repeatability of size for fish only to age 4. We estimated repeatability of body size using Generalized Linear Mixed Models as implemented in the R package *MCMCglmm* (Hadfield 2010, Wilson et al. 2010) with age as a fixed effect and fish ID as a random effect.

#### 2.4.1 Growth between sampling intervals

We used Generalized Additive Mixed Models (GAMMs) (Zuur et al. 2014) for each population separately to model variation in mean daily growth *G_d_* (in mm d^−1^) between sampling occasions using length *L*, *Age* (as categorical variable), *GDDs* over sampling intervals (by *Season* for populations sampled twice a year), and *D*_>0+_ as predictors, plus fish ID as a random effect. Since we expected potential non-linear relationships between the two predictors and *G_d_*, we used candidate smooth functions for *L* and *GDDs*. We carried out model fitting using the R package *mgcv* (Wood 2011) and model selection as in Section 2.3.1.

### 2.5 Recruitment

Marble trout spawn in November-December and offspring emerge in May-June. Females achieve sexual maturity when *L* > 200 mm, usually at age 3+ or older, and are functionally iteroparous (Meldgaard et al. 2007, Vincenzi et al. 2014a). We used density of fish with *L* > 200 mm as density of potential spawners at year *t* (*D*_s,t_). We used density of 0+ in September of year *t* as a measure of recruitment (*R*_t_). After pooling together population-specific data, we used Generalized Additive Models (GAMs) (Wood 2006) to model variation in *R*_t_ using *Stream*, density of potential spawners in September of year *t*-1 *D*_s,t-1_ and *GDDs* for year *t* up to emergence time (we assumed from January 1^st^ to May 31^st^ for standardization purposes) as predictors. We used candidate smooth functions for *GDDs* and *D*_s,t-1_ as we were expecting potential non-linear relationships between the two predictors and *R*_t_. Zak and Gac were not included, as they were sampled only in June of each year. We carried out model selection as in Section 2.3.1.

### 2.6 Survival

To characterize within-and among-population variation in survival and identify the determinants of variation, we modeled survival between sampling events for tagged fish and survival between age 0+ and 1+ for untagged fish.

#### 2.6.1 Survival of tagged individuals

Our goal was to investigate the effects of mean temperature, early density, floods, season, sex, age, sampling occasion, and growth potential on variation in probability of survival of tagged fish using continuous covariates (*D*_>0+_, mean temperature between sampling intervals 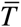, *Age,* and individual-level *k* and *L*_∞_ estimated using the growth model in Eq. (2.2) for growth potential) at the same time of categorical predictors (*Cohort* and *Time* for all populations, and *Season*, *Sex*, *Flood* when appropriate, as described hereafter). As only trout with *L* > 110 mm were tagged, capture histories were generated only for those fish. Full details of the survival analysis are presented in Appendix S1.

Two probabilities can be estimated from a capture history matrix: *φ*, the probability of apparent survival, and *p*, the probability that an individual is captured when alive (Thomson et al. 2009). In the following, for *φ* we will simply use the term probability of survival. We used the Cormack–Jolly–Seber (CJS) model as a starting point for the analyses (Thomson et al. 2009). The global starting model, that is the model with the maximum parameterization for categorical predictors, was different for each population. From the global model, recapture probability was modeled first. The recapture model with the lowest AIC was then used to model survival probabilities.

We modeled the seasonal effect (*Season*) as a simplification of full time variation, dividing each year in two periods: June to September (*Summer*) and September to June (*Winter*). Since the length of the two intervals (*Summer* and *Winter*) was different, we estimated probability of survival on a common annual scale, including *Season* as a potential predictor of probability of capture and survival in all populations that have been sampled twice a year. We included *Flood* (0 for no flood occurring during the sampling interval and 1 otherwise) as a potential predictor of probability of capture and survival for Lipo, Zak, Zadla, and Sve. For survival, we used *Flood* as a binary predictor to (a) test for differences in probability of annual survival of fish born before and after the flood, and (b) to estimate the decrease in probability of survival during a sampling interval in which a flood occurred with respect to sampling intervals with no occurrence of floods. We could estimate (a) only in Zak and Lipo, since there were not enough sampling occasions before the 2007 flood in Zadla and after the 2012 flood in Sve.

We used *Sex* as a predictor of survival in the populations of Zadla, Trebu, and Sve. Both *Age* and 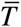 were introduced as either non-linear (as B-splines, Boor 2001) or linear predictors, while *D*_>0+_ only as a linear predictor.

To test the hypothesis of survival dependent on the growth potential of individuals, we introduced as potential predictors of survival the individual-level estimates of *L*_∞_ and *k*, as well as length at age 3+, *L*_3+_ (i.e., the typical age at sexual maturity), predicted using the vBGF with *L*_∞_ and *k* estimated at the individual level. We used the term *growth potential* as only some of the growth trajectories to the growth plateau were realized due to early mortality. Unless otherwise noted, probability of survival *φ* refers to an annual scale. We use *σ* to denote the probability of survival at the population level. We carried out the analysis of probability of survival using the package *marked* (Laake et al. 2013) for R.

#### 2.6.2 Survival from age 0+ to 1+

Because fish were not tagged when smaller than 110 mm (thus 0+ fish were not tagged), we assumed a binomial process for estimating the probability *σ*_0+_ of first overwinter survival (0+ to 1+) (see Appendix S2 for details on the estimation of *σ*_0+_). After pooling population-specific data, we tested for density-dependent survival σ_0+_ by estimating a linear model with *Stream*, *D*_>0+_ at year *t* in September, and their interaction as predictors of the estimate of 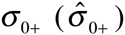. Following Vincenzi *et al.* (2008b, 2010b), we log-transformed both 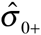 (adding 0.01 to each value as some values of 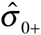 were equal to 0) and 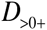. We excluded the years following flood events from the analysis. We carried out model selection as in Section 2.3.1.

### 2.7 Growth-survival trade-off

To test the trade-off between expected growth and expected probability of survival across marble trout populations, we used estimates of growth and survival that represent “normal” behavior. For each population we used population-level estimates of *k*, *L*_∞_, and probability of survival for cohorts that were born before flood events (i.e. estimates of growth model in Eq. 2.2. with *Coh*_fl_ as predictor of vBGF’s parameters), based on results from Vincenzi *et al.* (2008a,b, 2014b). In addition, for Zak and Gac we used estimates for cohorts other than the parental (i.e. estimates with *Coh*_p.fl_ as predictor for Zak and *Coh*_p_ for Gac).

In order to account for uncertainty in the estimation of vBGF parameters when testing the relationship between growth and survival rates, we carried out Monte Carlo simulations to obtain a distribution of *L*_∞_ − *σ* and *k* − *σ* correlation estimates (***r***), and of probability of *r* being different from zero (***p***) (Appendix S3).

## 3 Results

### 3.1 Density and Floods

In all streams, population density varied through time, with the highest Coefficient of Variation (CV) of *D*_>0+_ observed for Zak (0.74) and the lowest for Trebu (0.21). The CV of *D*_>0+_ was not correlated with number of sampling occasions (Pearson’s *r* = 0.14, *p* = 0.70). The populations with higher CV of *D*_>0+_ experienced at least one severe flood event during the study period (Fig. 2), thus indicating a major role of flood events in increasing fluctuations in density. There was a strong and significant lagged correlation (lag of 1 year) between *D*_0+_ and *D*_>0+_ in September in all populations (*r* between 0.55 [Stu] and 0.80 [Sve]), except for Huda (*r* = 0.38, *p* > 0.05), which had instead a significant positive lagged correlation (after 3 years) between *D*_0+_ and density of potential spawners *D*_s,t_ (*r* = 0.64). This result indicates that recruitment was largely driving variation in population density.

**Fig. 2.**
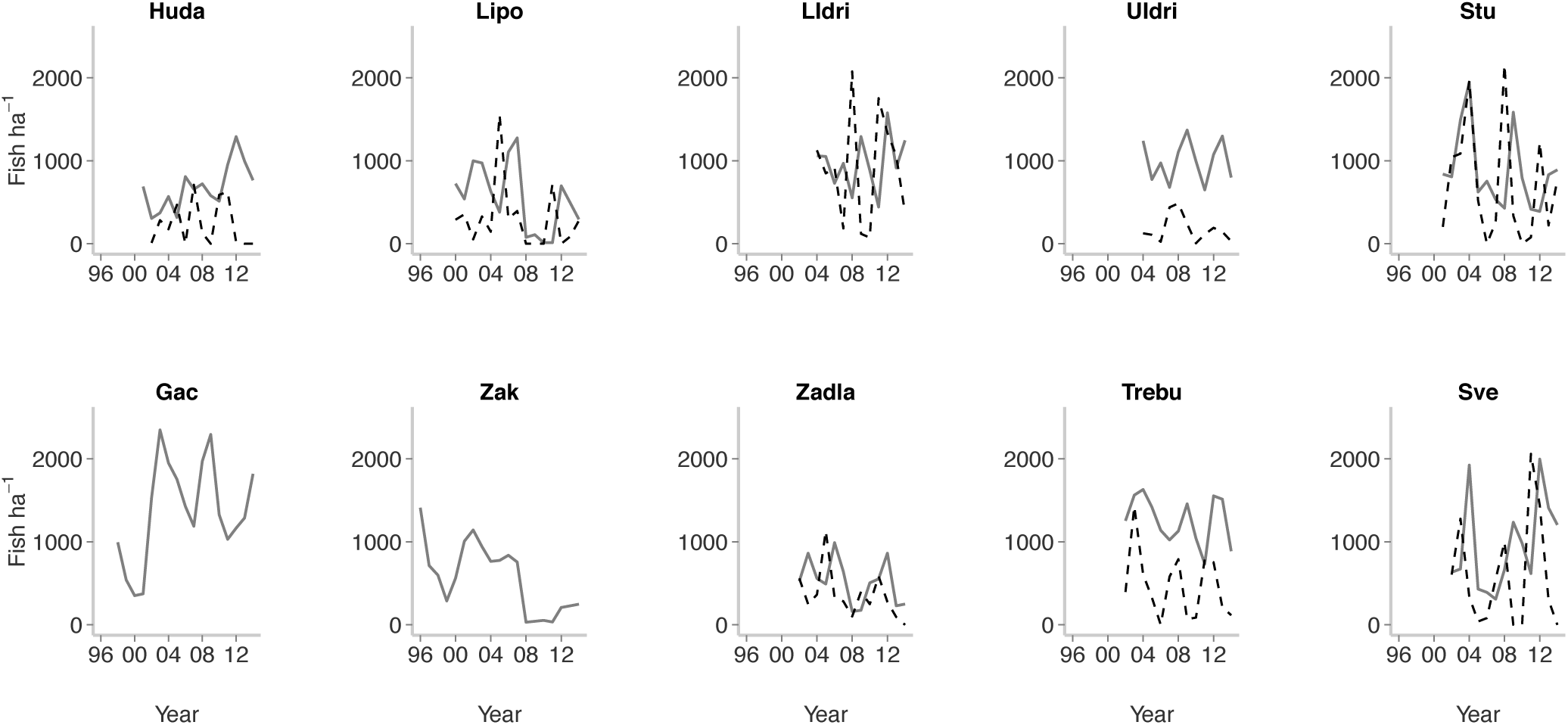
Density over time of age-0+ marble trout (dashed line) and older than age-0+ (solid line) in September for all populations excluding Zak and Gac (sampling in June and density of age-0 not estimated). Severe flash floods and debris flows occurred in Zadla (2007 and 2012), Zak (2007), Lipo (2004, 2007 and 2009), Sve (2012), Stu (2004), Sve (2004).

### 3.2 Growth and body size

#### 3.2.1 Size at age 0+

Mean length of age 0+ fish 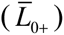 in September was density-dependent after accounting for the effect of *Stream*, with the populations of Huda, Sve and Trebu showing smaller 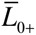 -at-density than the other populations (Table S3 and Fig. 3). Mean condition factor at age 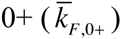 was between 0.87 (Huda, 2002]) and 1.07 (Lipo, 2012). As found for 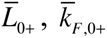 was density-dependent after accounting for the effect of *Stream* (Table S3). 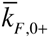 was positively correlated with 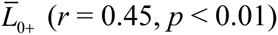, thus larger fish were on average in better condition. The populations of Huda, Sve and Lipo had the lowest 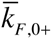 -at-density.

**Fig. 3.**
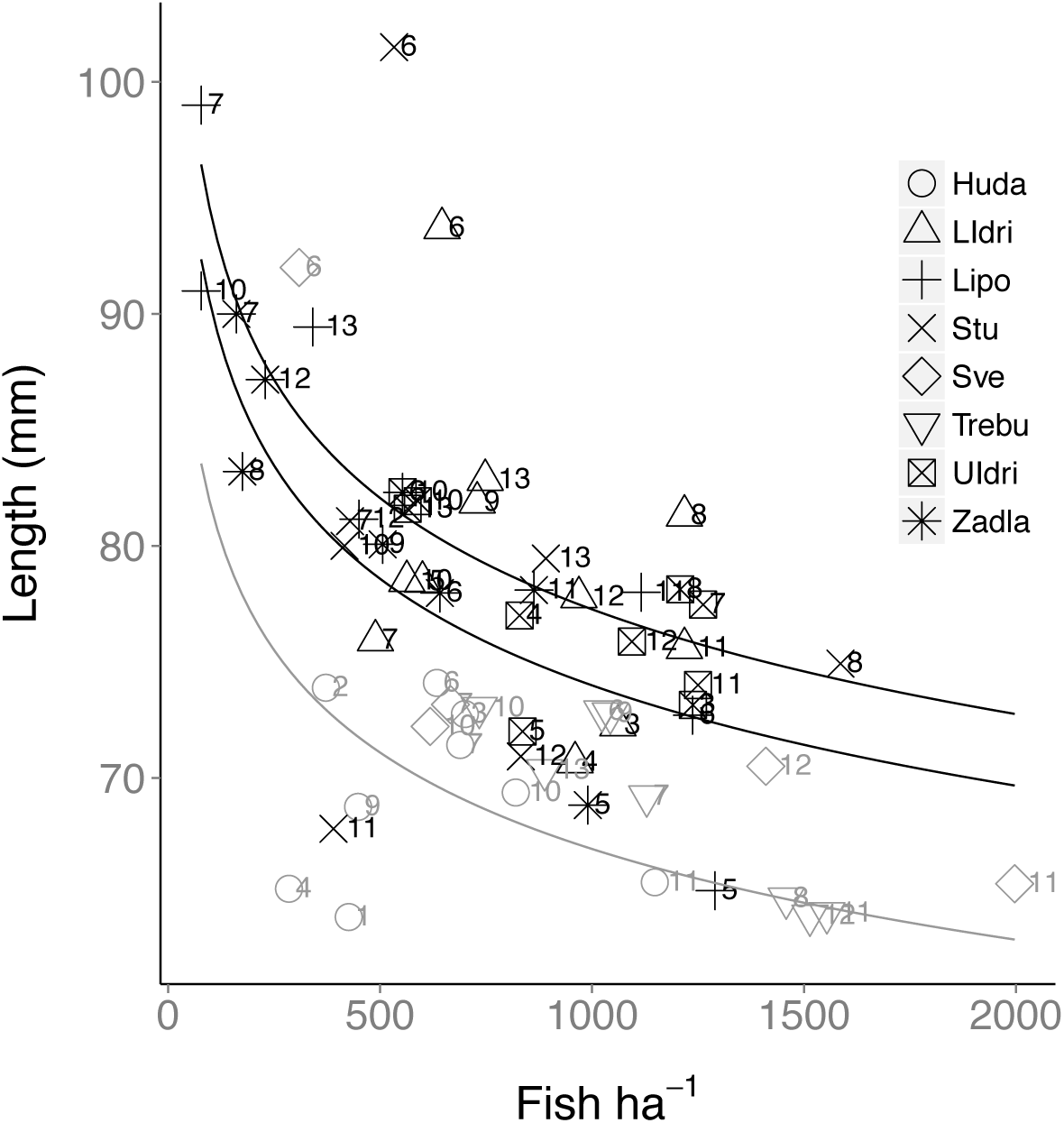
Density-dependent mean body size of cohorts at age 0+ in September. Numbers indicate the cohort (e.g., 7 = fish born in spring of 2007). The gray line is the regression line (on a log-log scale) for Huda, Sve, Trebu, while the black lines bound the regression lines for the other populations (see Table S1).

#### 3.2.2 Lifetime growth trajectories

Empirical growth trajectories showed substantial individual variation in growth rates and size-at-age (Fig. 4), thus supporting the choice of a growth model with individual random effects. Using the complete datasets (Data_W_) in the vBGF model with no predictors for vBGF’s parameters, asymptotic size *L*_∞_ and growth coefficient *k* were strongly and positively correlated in all populations (Pearson’s *r* from 0.70 [Zak] to 0.97 [UIdri and Zadla], *p* < 0.01 in all populations). Mean growth trajectories predicted by the vBGF model with no predictors tended to plateau rapidly after sexual maturity in Huda, Trebu, Zak, and Gac (Fig. 4). The best growth model for every population had *Cohort* as a predictor of either *L*_∞_ or *k* or both (Table S4).

**Fig. 4.**
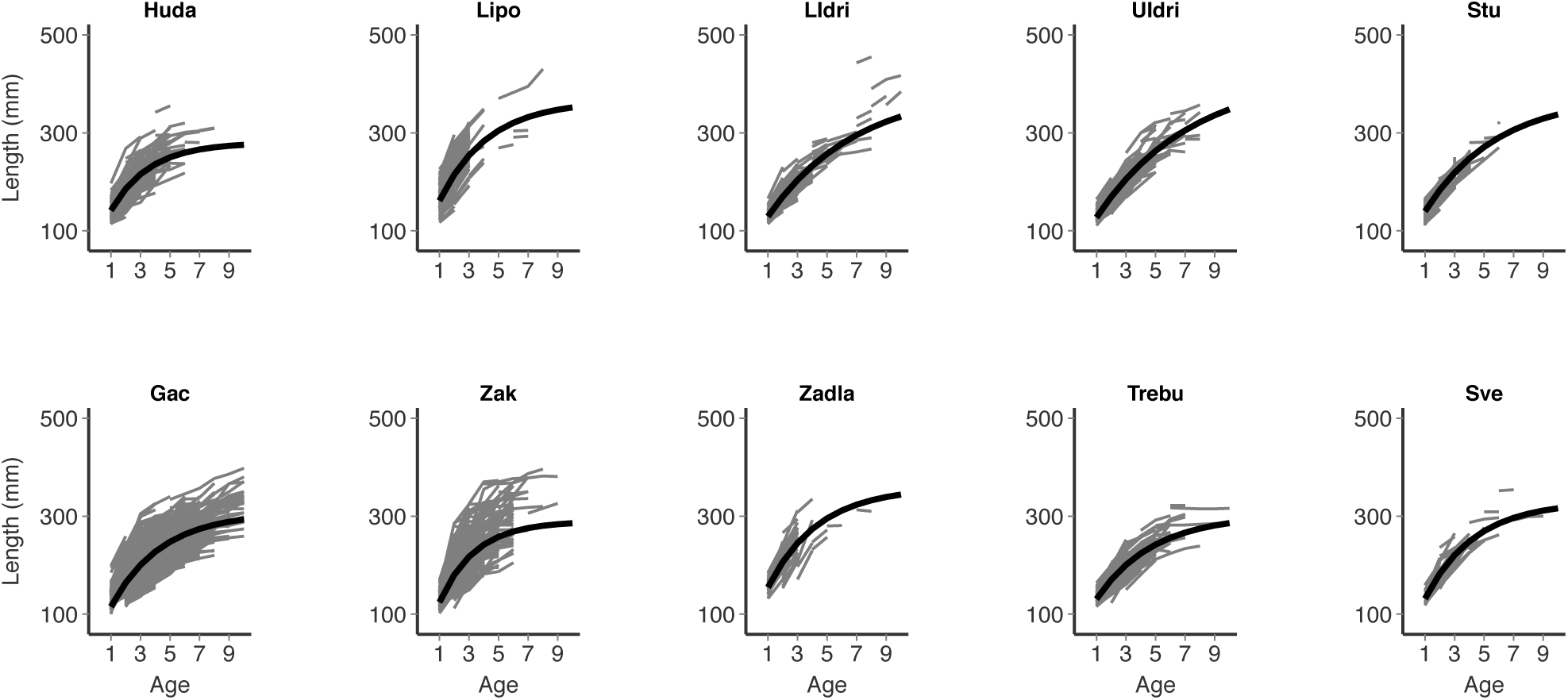
Growth trajectories of marble trout that were sampled at least two times, in June (Zak and Gac) or in September (all other populations). Parental cohorts of Zak and Gac are not included. Thick lines are the predictions of the vBGF models for the average individual in the populations (i.e., no predictors for either *L*_∞_ and *k* and individual random effects set to 0).

In streams affected by flood events, cohorts born after flood events grew faster and had greater asymptotic size than cohorts born before the flood (Fig. 5), thus pointing to a substantial acceleration of growth following episodes of massive mortality.

**Fig. 5.**
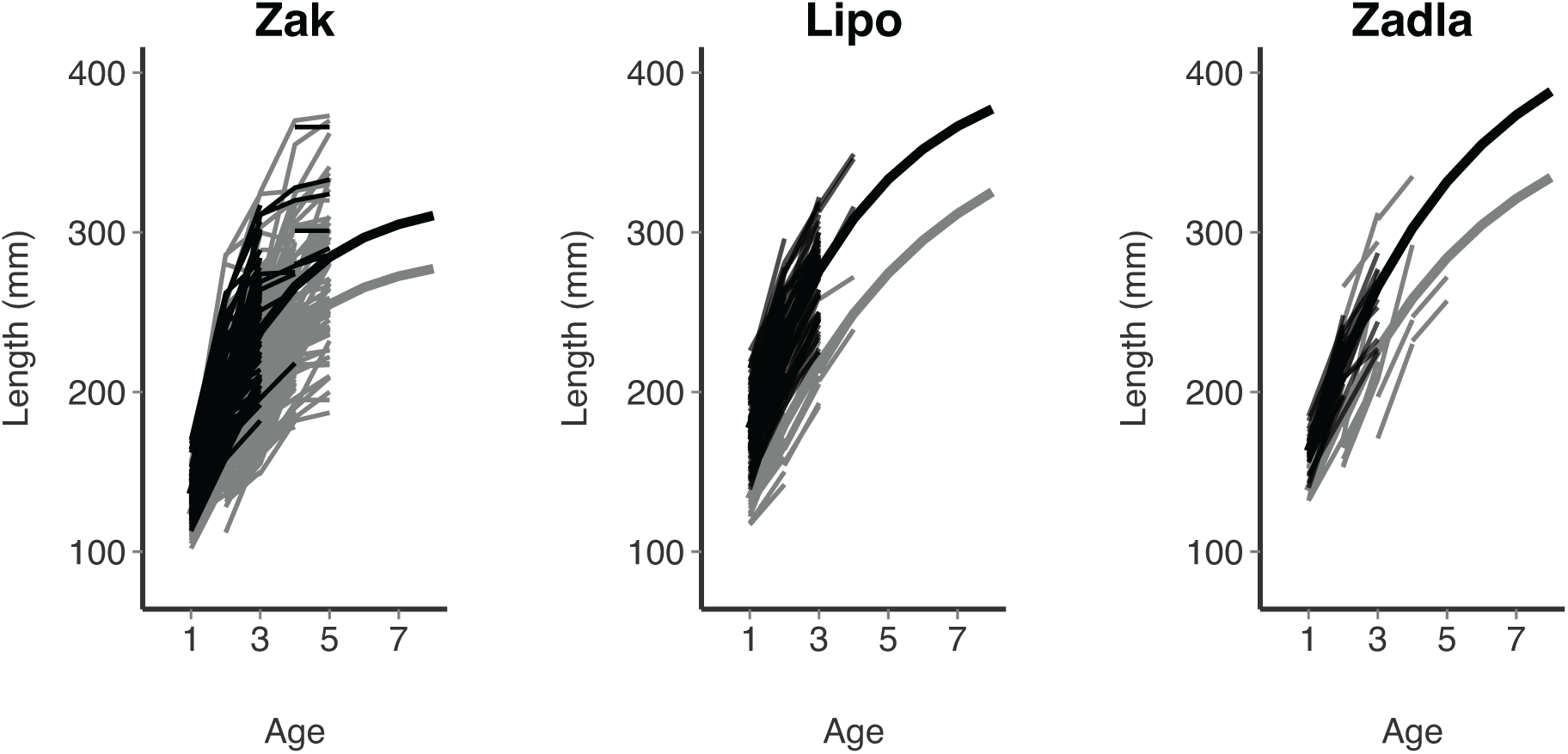
Cohorts born after floods (black lines) grew faster and had higher estimated asymptotic size than cohorts born before the flood event (gray lines). Thick black and gray lines are the predicted mean growth trajectories for cohorts born after and before the flood, respectively. Mean[95%CI] - Zak: Pre-Flood, *L*_∞_ = 286.40 mm [279.89-292.91], *k* = 0.41 y^−1^ [0.39-0.43], t_0_ = -0.36 y; Post-Flood, *L*_∞_ = 321.65 mm [305.40-337.90], *k* = 0.40 y^−1^ [0.36-0.44], t_0_ = -0.36 y [-0.41-(-0.32)]. Lipo: Pre-Flood, *L*_∞_ = 384.79 mm [331.82-437.76], *k* = 0.21 y^−1^ [0.15-0.26], t_0_ = -1.06 y [-1.29-(-0.83)]; Post-Flood, *L*_∞_ = 411.94 mm [360.04-463.84], *k* = 0.27 y^−1^ [0.20-0.35], t0 = -1.0556 y [-1.29-(-0.83)]. Zadla: Pre-Flood, *L*_∞_ = 388.88 mm [323.40-454.36], *k* = 0.22 y^−1^ [0.14-0.30], t_0_ = -1.01 y [-1.40-(-0.61)]; Post-Flood, *L*_∞_ = 446.48 mm [291.30-601.67], *k* = 0.23 y^−1^ [0.09-0.37], t_0_ = -1.007 y [-1.40-(-0.61)].

When using Data_D_ (i.e., also including *GDDs* and *D*_>0+,_*_born_* as predictors of *k* and *L*_∞_), the best models were the same as those found when using Data_W_ (Table S4). However, for every population, models with *D*_>0+_ and/or *GDDs* as predictor of either *L*_∞_ or *k* or both were better than models with no predictors for both *L*_∞_ and *k*, thus pointing to a potential role played by early density and temperature in determining growth trajectories. The effects of the two variables were on opposite directions. For every population, *L*_∞_ tended to get smaller with increasing *D*_>0+_. For every population except Huda and Trebu, *L*_∞_tended to get larger with *GDDs* in the first year of life (see Online Code and Results).

Sex-specific growth trajectories differed only in Trebu, although predicted length-at-age started to differ only after age 2+. In Zadla, Sve, and Zak (using dataset DataS for the latter), sex-specific 95% CIs of *k* and *L*_∞_ largely overlapped, indicating no difference in the mean growth trajectories of males and females (see Online Code and Results).

#### 3.2.3 Growth between sampling intervals

For each population, at least one of the best models included density as a predictor of mean daily growth *G_d_*, although the density coefficient was not always negative. *G_d_* was well predicted by the best growth models, with coefficient of determination *R*^2^ ranging from 0.39 (Huda) to 0.70 (Svenica) (Table S5 and Fig. S3). Among the populations sampled once a year in September, marble trout in Zadla had greater mean daily growth than fish in Sve and Trebu. Although *GDDs* were always retained as a predictor in the best models, their effect on growth varied with *Season* and *Stream*. A positive effect of *GDDs* on growth was observed for *Winter* growth in LIdri, UIdri, and Lipo and for *Summer* growth in Lipo (Table S5).

#### 3.2.4 Repeatability

We obtained very high estimates of repeatability of body size measurements throughout lifetime for all populations (mean estimates from 0.7 [Sve] to 0.88 [Lipo]). In all populations except Zadla (only 9 fish were sampled at age 1+ and age 3+ in September), we found a strong positive correlation between size at age 1+ and size at age 3+ (Pearson’s *r* from 0.49 [Gac] to 0.86 [Sve], *p* < 0.01 for all populations). Along with the positive correlation found between asymptotic size *L*_∞_ and growth coefficient *k*, the high repeatability of body size points to the maintenance of size ranks through marble trout lifetime.

### 3.3 Recruitment

The best model of recruitment *R*_t_ had *Stream*, density of spawners *D*_s,t-1_ and *GDDs* as predictors, and explained ~28% of the variation in recruitment (Table S6). The continuous predictors had opposite effects on recruitment: *D*_s,t-1_ had a linear and positive effect on *R*_t_, while the effect of *GDDs* on *R*_t_ was linear and negative. A model with *GDDs* not included as a predictor had only slightly less support than the best model (Akaike weight of the best model and the model without *GDDs* were 0.57 and 0.43, respectively).

### 3.4 Survival

#### 3.4.1 Survival of tagged individuals

In Tables 1 and S7 we show the best capture and survival models for each population, respectively. Probability of capture using the best capture model was high in every population (from 0.7 [Huda] to 0.88 [Zak]) and tended to be higher for populations sampled once a year.

**Table 1.**
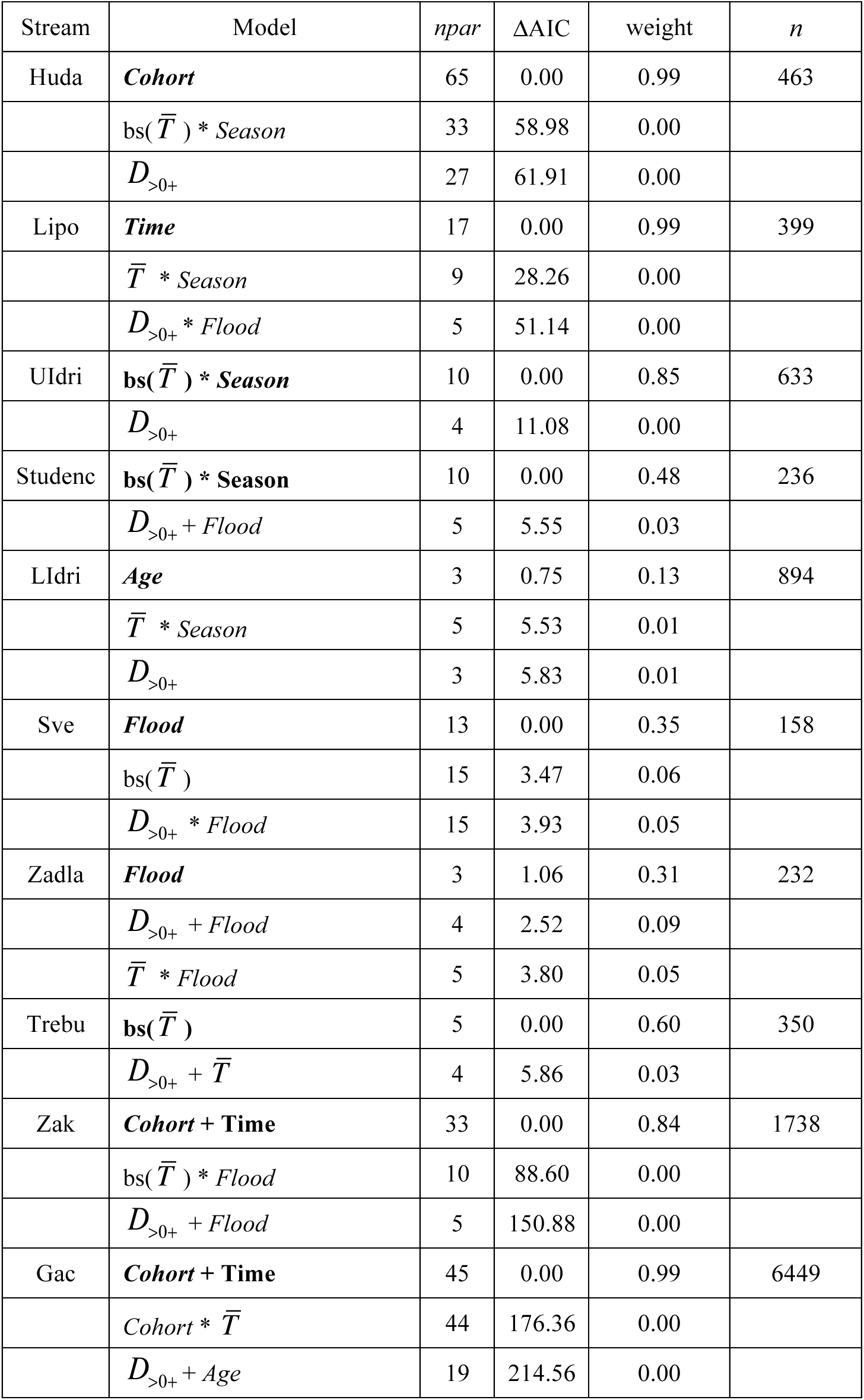
Best models of probability of survival using the population-specific probability of capture in Table S4. For each population, we report (in **bold**) the model with the least number of parameters among those within 2 points of AIC from the best model (ΔAIC = 0 if it is the model with the smallest AIC among all fitted models), the best model with mean temperature between sampling occasions 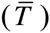 as predictor, and the best model with *D*_>0+_ as predictor. *Time* = interval between two consecutive sampling occasions. In case the best model with 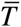 as predictor is the best overall model, we only report two models of survival for that population. *bs* means that the relationship temperature and probability of survival has been modeled as a B-spline function. *npar* = number of parameters of the survival model; *n* = number of unique tagged fish. See Online Code and Results for a complete list of models for each stream.

##### Effects of Density and Temperature

There was no evidence of density-dependent survival of tagged fish in any of the populations. Models with 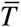 as a predictor of probability of survival *φ* were the best-supported models in UIdri, Stu, and Trebu (Table 1). In each of those models, the relationship between mean annual (Trebu) or seasonal (UIdri, Stu) temperature and *φ* was modeled with B-splines. However, a more fine-grained analysis of the relationship between 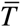 and *φ* showed that the inferred relationships are likely to be a statistical artifact, as they were not biologically interpretable (Fig. S4).

##### Effects of Season, Sex, Flood, and growth potential

There were no substantial seasonal differences in *φ* in any of the populations that were sampled twice a year, except for Lipo (Table 2), in which June-to-September (*Summer*) *φ* was greater than September-to-June (*Winter*) *φ* due to the high survival rates in the summers following floods. There were also not any substantial differences between *φ* of males and females in the three populations with sex data, i.e., Sve, Zadla, or Trebu (Table 2).

**Table 2.**
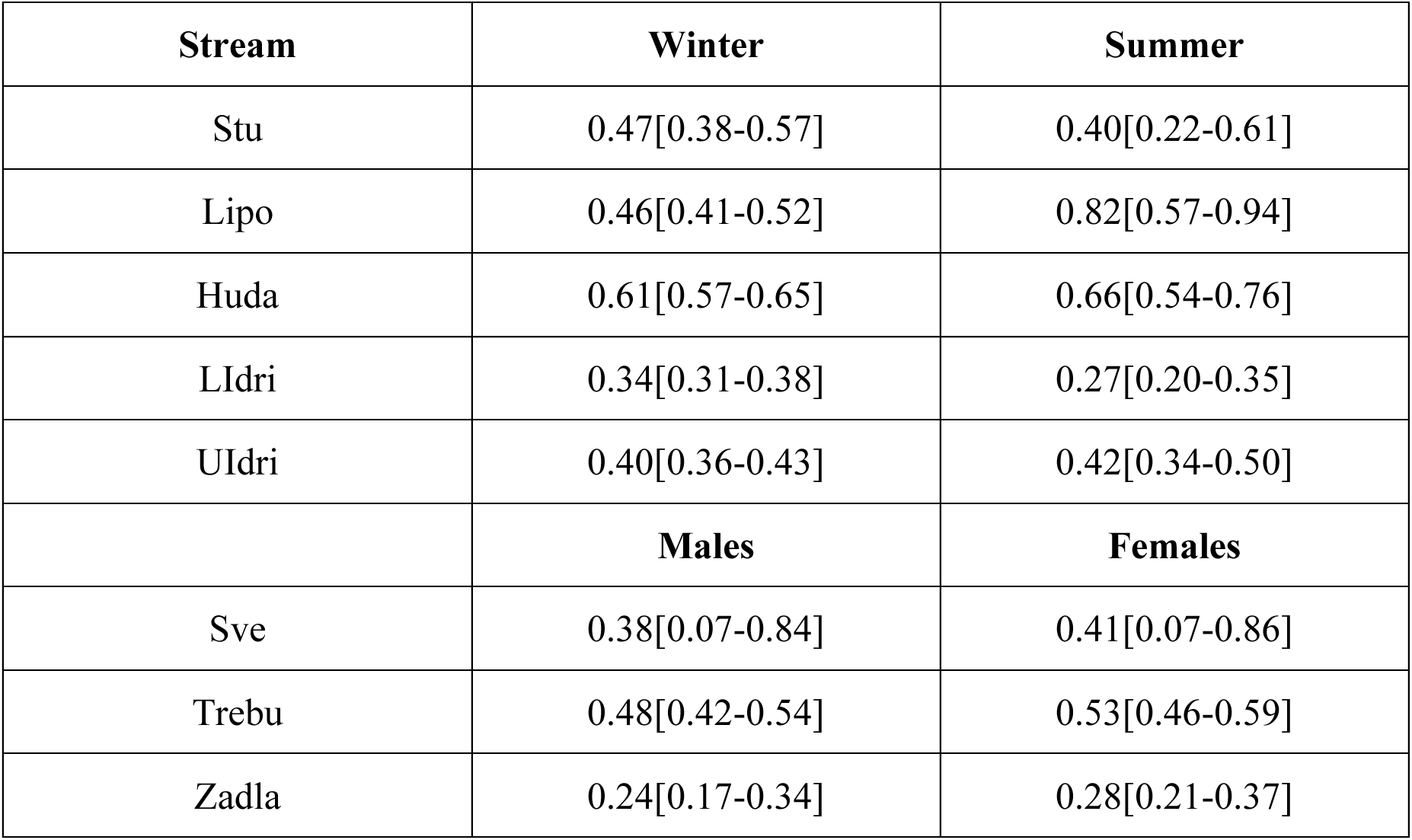
Mean [95% CI] of (a) September-to-June (*Winter*) and June-to-September (*Summer*) probability of survival (on a common annual scale) for the marble populations that have been sampled twice a year, (b) sex-specific probability of survival in the populations of Sve, Trebu, and Zadla.

Models with *Age* as a predictor of probability of survival had very little support for all populations except for LIdri, in which the best model had *Age* as the only predictor of survival (Table 3). However, the higher probability of survival at older ages was mostly driven by a few fish that were sampled in LIdri when older than age 12 (Fig. S5).

For Zadla and Sve, the best models had *Flood* as a predictor of *φ*, while for Lipo and Zak models with *Time* better explained variation in survival (Table 1). Floods caused a > 80% reduction in the probability of survival in Zak, Sve, and Lipo and a ~ 55% reduction in Zadla relative to sampling intervals not affected by floods (Fig. 6). In both Zak and Lipo, the probability of survival of fish born after the flood was noticeably greater than that of fish born before the flood, although the confidence intervals of the estimates for Lipo overlapped (Fig. 6).

**Fig. 6.**
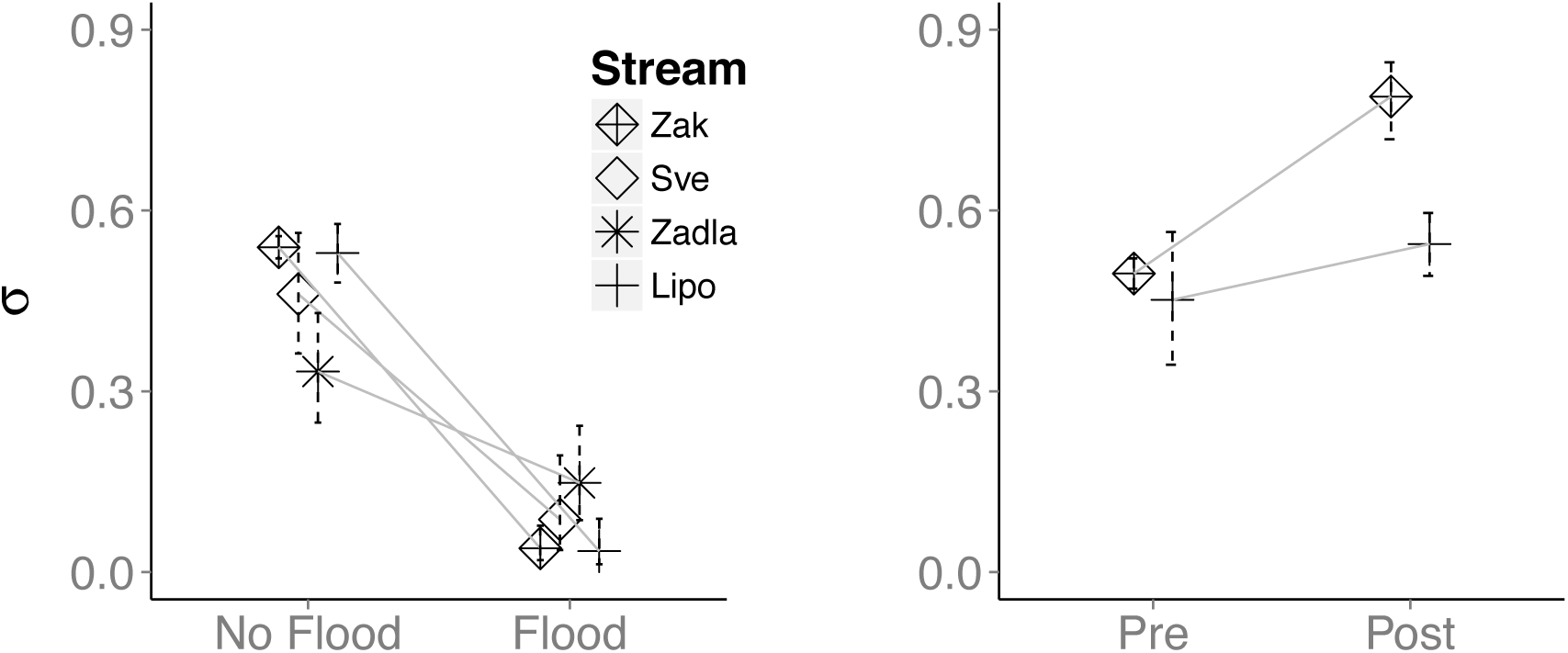
(a) Annual probability of survival (mean and 95% confidence intervals) in years with no flood and in years with flood(s) for the populations of Zak, Sve, Zadla, Lipo. (b) Annual probability of survival (mean and 95% confidence intervals) - in years with no floods - of cohorts born before or after the flood event in the populations of Zak and Lipo.

There was very little support for models of *φ* as a function of fish growth potential in any of the populations: models including either *L*_∞_, *k*, or predicted *L*_3+_ lacked support in every population and no clear relationship with *φ* was observed for any of those potential predictors.

#### 3.4.2 Survival from age 0+ to 1+

The probability of early survival *σ*_0+_ was density-dependent (Fig. 7), although the model explained only ~15% of the variation in early survival. The best model for *σ*_0+_ had only density as a predictor, with no differences among populations in either the intercept or the slope of the regression. The probability of survival over the first winter was typically lower than survival at older life stages (Fig. 7 and Fig. 8).

**Fig. 7.**
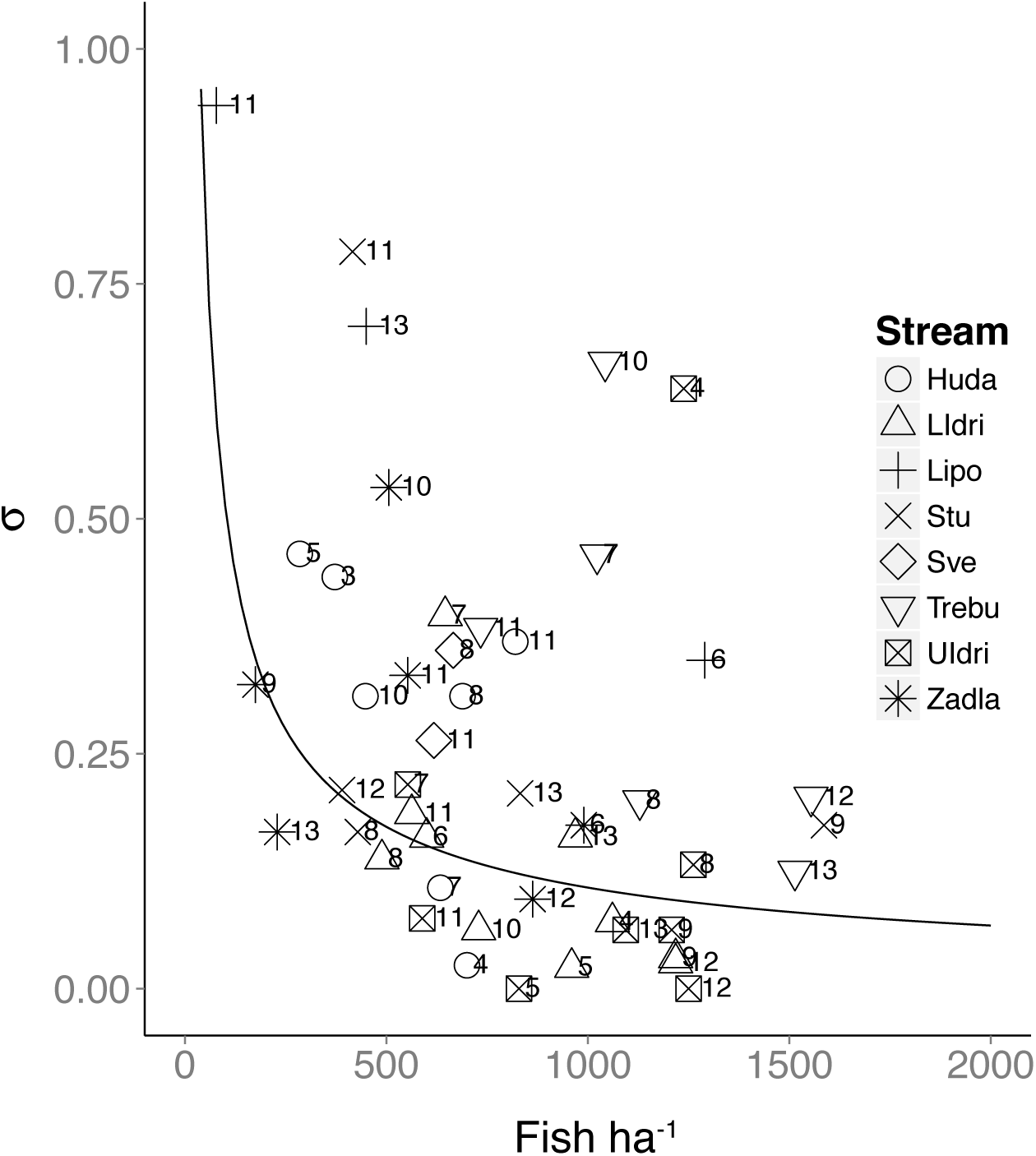
Density-dependent survival from age 0+ in September to age 1+ in June (Huda, Lipo, LIdri, UIdri, Stu) or September (Sve, Zadla, Trebu). Survival is estimated on a common annual scale. The regression line is for the model 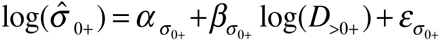, 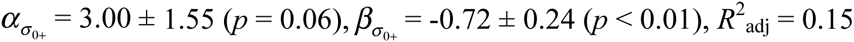.

**Fig. 8.**
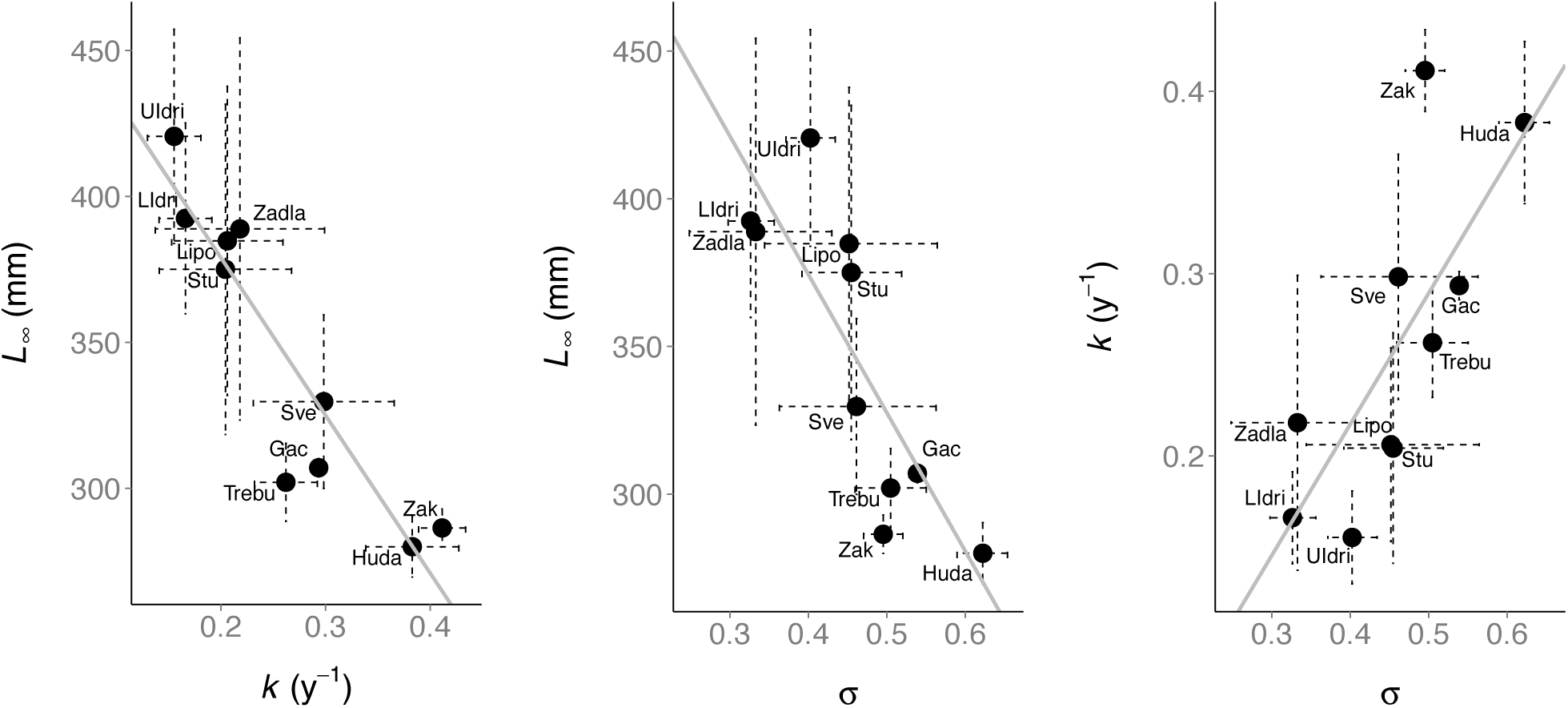
(a) Relationship between *k* and *L*_∞_ estimated at the population level. For Zak, Zadla, Lipo we used the parameters estimated for cohorts born before the flood (pooled together) and different from the parental cohort (for Zak and Gac). Relationship between (b) *L*_∞_ and probability of annual survival *σ*, and (c) *k* and *σ*. For Zak, Zadla, Lipo, and Stu we used *σ* estimated for years or sampling intervals in which a flood did not occur and for fish born before the flood and different from the parental cohort (the latter for Zak and Gac). Estimates of regression parameters are reported in Online Code and Results.

### 3.5 Growth-survival trade-off

We found strong correlations between population-level *L*_∞_ and *k* (*r* = -0.92, *p* < 0.01), *L*_∞_ and σ (*r* = -0.82, *p* < 0.01), and *k* and *σ*(*r* = 0.74, *p* < 0.01) (Fig. 8). The strength of the relationships did not substantially change when the newly created populations of Zak and Gac were excluded (*L*_∞_ and *k*: *r* = -0.92, *p* < 0.01; *L*_∞_ and *σ*: *r* = -0.83, *p* = 0.01; *k* and *σ*: *r* = 0.83, *p* = 0.01). We found a strong negative correlation between asymptotic length and the growth coefficient of the vBGF estimated at the population level, indicating that the average predicted growth trajectories often cross through marble trout lifetime (Fig. S6). Furthermore, the mean condition factor at age 1+ in September was positively correlated with the point estimate of *L*_∞_ at the population level (Fig. S7).

Using Monte Carlo simulations, Pearson’s *r* for *L*_∞_ and *σ* was 0.74 (95% CI: [-0.89-(-0.48)]) (16% of *p* values were greater than 0.05), and 0.68 [0.46-0.84] for *k* and *σ* (28% of *p* values were greater than 0.05). These results indicate that marble trout of UIdri, LIdri, Zadla (and potentially Lipo and Stu) are characterized by a faster pace of life with faster growth, higher condition factor, and shorter lifespan.

## 4 Discussion

We discuss our results on the causes of variation in growth, survival, and recruitment both within and among marble trout populations, how those results help advance our understanding of demography and life-history processes in animal populations, the pieces of missing information that can further advance our understanding of those processes, and the implications of our findings for the conservation of marble trout.

In all populations, the effects of population density on vital rates were generally limited to the early stages of life and individual growth trajectories appeared to be established in the first year of life. Growing degree-days early in life may influence lifetime growth trajectories, but there were no evident effects of mean temperature or growing degree-days on survival and recruitment. Population density naturally varied over time in all populations, with occasional flash floods and associated debris flows causing massive mortalities and threatening the survival of the affected populations. Apart from flood events, variation in population density within streams was largely determined by variation in recruitment, with survival of older fish being relatively constant over time within populations, but substantially different among populations. A fast-to slow-continuum of life histories in marble trout populations seems to emerge, with slow growth associated with higher survival at the population level, possibly determined by food conditions and size-dependent age at maturity.

### 4.1 Growth

The main findings of our analysis of marble trout growth are: (1) density-dependent early growth, which is found commonly in salmonids (Jenkins et al. 1999, Vincenzi et al. 2012b); (2) lifetime growth trajectories that vary substantially by population and year-of-birth cohort; and (3) maintenance of size ranks through lifetime with a high correlation between size-at-age early and later in life, indicating an early determination of lifetime growth trajectories and a smaller role of population density and temperature experienced throughout the lifetime after the first year of life.

In each population, the best model of lifetime growth trajectories included cohort as a categorical variable for either the asymptotic size or the growth coefficient of the von Bertalanffy model for growth or both, indicating that the resulting predicted growth trajectories of fish are more similar to those of other fish in the same cohort than to those of the population as a whole. These early, environment-induced modifications of the phenotype may reflect either constraints or adaptations and are often ascribed to climatic vagaries during early development giving rise to cohort effects (Lindström 1999; Mangel 2008; Monaghan 2008). Population density may also have an effect on growth, in particular due to the effect of density on food availability (Vincenzi et al. 2008b), or the occupation of spaces with low profitability at higher density (Newman 1993). Models including population density in the first year of life and/or growing degree-days performed distinctly worse than the best model, but better than the model with no predictors. This may suggest that either or both population density and growing degree-days explain part of the variability in lifetime growth trajectories of marble trout, although further studies in controlled environments that can elucidate the role of temperature and density are needed.

Population density and growing degree-days between sampling occasions were often included in the best models of individual growth between sampling occasions, but their effect sizes were small and their direction not consistent across populations. Size ranks tended to be maintained throughout lifetime: individuals that were relatively large early in life tended to remain larger than their conspecifics throughout their lifetime. High heritability of growth (Carlson and Seamons 2008), maternal decisions on the timing and location of spawning (Letcher et al. 2011), dominance, possibly determined by higher metabolic rates giving easier access to resources (Gilmour et al. 2005, Reid et al. 2011), are all processes that may explain the maintenance of size ranks throughout marble trout lifetime.

### 4.2 Effects of floods

Flash floods occurred in Zadla, Lipo, Sve, and Zak and caused heavy losses, but the populations were able to rebound rapidly to pre-flood density. Some studies on fish populations affected by floods have shown that adult and juvenile trout displaced downstream have returned upstream after the flood recession, and that re-colonization by unaffected sub-populations living nearby can also be quick (Dedual and Jowett 1999, Ortlepp and Murle 2003, Weese et al. 2011). However, Zadla, Sve and Zak are all fragmented streams in which opportunities for upstream movement and re-colonization after fish have been flushed downstream by fast waters are severely limited - if not impossible - due to the presence of large and impassable waterfalls. On the other hand, our results show that a few survivors can rapidly increase population size to pre-flood levels due to intrinsic processes, in particular the relaxation of density-dependent limitations on growth and survival (Lamberti et al. 1991, Vincenzi et al. 2012a, 2014a, Ohlberger and Langangen 2015). Both processes emerged as clearly important ones in our analyses and led to swift recoveries after the population collapses induced by the flood events.

### 4.3 Recruitment

Recruitment is the most variable and influential vital rate for many fish populations (Ricker 1975), especially for short-lived species (Bjørkvoll et al. 2012). Recruitment in marble trout was highly variable and marble trout populations were recruitment-driven, as indicated by the strong one-year lagged correlation between density of newborns and density of older-than-newborn trout in all remnant populations except Huda. Although variable recruitment may lead to increased uncertainty in recovery time after a population collapse (Kuparinen et al. 2014), the consistent production of strong cohorts after population collapses in marble trout enhances population recovery (see also Vincenzi *et al.* 2014a).

The relative balance between spawning stock size (i.e. the number or density of spawners) and environmental factors as determining recruitment in riverine salmonids is still debated and probably context-specific (Einum 2005, Lobón-Cerviá 2005, Nicola et al. 2008). In marble trout, recruitment linearly increased with the density of potential spawners, but neither density of potential spawners nor growing degree-days explained a large part of the observed variability in recruitment. These results may indicate that there are other un-studied, but potentially important, environmental factors (e.g. water flow, dissolved oxygen) for marble trout recruitment (Nicola et al. 2009, Unfer et al. 2011). On the other hand, in salmonids only a fraction of potential spawners have reproductive success, and the productivity of successful spawners is highly skewed (Esteve 2005). The density of potential spawners estimated according to size is thus only a crude proxy of the density of real spawners or distribution of reproductive success among spawners. By helping us identify the successfully reproducing individuals, molecular pedigree reconstruction (Anderson and Garza 2006, Kruuk and Hill 2008, Pemberton 2008) will greatly advance our understanding on the determinants of recruitment in marble trout.

### 4.4 Survival

Density-dependent early survival has been often found in salmonids (Jonsson and Jonsson 2011) and other fish species (Minto et al. 2008), although there are examples of salmonid populations showing density-dependent survival only at the adult stage (Lobón-Cerviá 2012) or constant loss rates (Elliott 1989). In marble trout, the probability of survival early in life was density-dependent and typically lower than (although comparable to) survival at older life stages, as previously observed for other salmonids (Schlosser 1995).

Survival of tagged fish between sampling occasions showed little variability within populations, except after the occurrence of floods, thus showing that population regulation occurred at early life stages and that environmental conditions within streams were relatively stable. In the populations with the highest number of fish tagged (Gac, Zak, Huda), the best models included year-of-birth cohort as a predictor of survival. This result indicates that cohort effects are strong determinants of growth, survival, recruitment, and thus of population dynamics of marble trout (Lindstrom and Kokko 2002). We found no evident effect of individual growth potential on probability of survival, which revealed that - within populations - there were no trade-offs between growth, or traits associated with growth (e.g. metabolism), and survival.

### 4.5 Growth-survival trade-off

We found a strong correlation between point estimates of parameters of the vBGF and point estimates of survival at the population level. The Monte Carlo simulations and other sensitivity analyses (Appendix S4) indicate that this result was quite robust to uncertainty in parameter estimation and possible confounders. vBGF’s parameter estimates can seldom be interpreted separately, especially when only a few older fish are measured (Vincenzi et al. 2014b). Asymptotic size is the parameter of the vBGF with the most immediate biological interpretation and we thus focus on asymptotic size in our discussion of the growth-survival trade-off at the population level. Marble trout living in Huda, Trebu, and Sve had a lower mean estimated asymptotic size and also smaller mean size at age 0+. It is also evident from the mean growth trajectories predicted by the model that growth tended to plateau rapidly after sexual maturity in Huda, Trebu, Zak, and Gac, while growth did not slow down as quickly in populations with higher mean estimated asymptotic size, such as UIdri, Lidri, and Zadla.

One hypothesis to explain the growth-survival trade-off at the population level is differences in the prevalence of cannibalism, since higher cannibalism would sustain growth at the expense of the mean probability of survival in the population (Mangel and Abrahams 2001, Finstad et al. 2006). However, marble trout living in Huda, Trebu, and Sve were also smaller when 0+, a result that does not support the cannibalism hypothesis, as 0+ are unlikely to eat other fish due to their small size and food preferences. Another hypothesis is alternative genetically determined life-history strategies, with some populations adopting faster life histories through a preferential allocation of resources to growth (Mangel and Stamps 2001). However, Zak and Zadla are genetically equivalent (Zak was created with progeny of wild-caught Zadla trout) and are at the opposite corners of the trade-off surface.

Variation in growth and survival across populations thus seemed to be mostly determined by environmental conditions - trophic conditions in particular, since water temperature is generally well within the optimal range for salmonids (Elliott and Elliott 2010) - and size-dependent age at maturity. In salmonids and other fish species, faster growth early in life can accelerate sexual maturity (Alm 1959, Craig 1985, Jonsson et al. 2013) and increase mortality of spawners due to energetic limitations (Berg et al. 1998). The hypothesis of mean growth largely determined by food conditions is also supported by a positive correlation between asymptotic size and mean fish condition across populations, along with evidence of variation in growth along the same stream. In all streams, we found consistently bigger size-at-age of fish occupying the uppermost part of the stream – where a larger portion of stream drift is available since no fish are living upstream - than of those fish living further downstream (Vincenzi et al. 2010b, 2014b). The only exception was LIdri, in which the biggest fish were found in a big pool located in the downstream portion of the sampling area.

Marble trout of UIdri, LIdri, Zadla (and potentially Lipo and Stu) were characterized by a faster pace of life with faster growth, higher condition factor, and shorter lifespan. A similar fast to slow continuum of life histories has been found in seabirds (e.g., kittiwake *Rissa tridactyla*) and marine species (e.g., leatherback sea turtle *Dermochelys coriacea*) living in the Atlantic (faster life histories) and Pacific Ocean (slower life histories) (Suryan et al. 2009), and on smaller geographical scale in slimy sculpin *Cottus cognatus* (Bond et al. 2015) between natural and regulated rivers. Differences in sculpin life-history traits within and among rivers closely followed spatial patterns in food availability, with faster growth, higher condition factor, and lower survival in food-rich regulated rivers than in natural rivers. Within regulated rivers, sculpin at sampling sites near dams (where more food was available) grew more rapidly and matured earlier than fish at sites farther downstream. Increased growth in the regulated rivers likely corresponded with earlier onset of sexual maturity and thus higher mortality in those populations (Bond et al. 2015).

### 4.6 Implications for conservation and future work

Small, fragmented populations are usually assumed to be particularly vulnerable to extinction, since they may be strongly affected by environmental and anthropogenic sources of disturbance that can cause population size to fluctuate greatly and possibly drop to very low densities (Lande 1993). Our work confirms that the major risk of extinction for marble trout is represented by extreme events (Vincenzi et al. 2008d) and that population sizes in the absence of floods are generally large enough to be safe from the effects of stochastic fluctuations. In addition, occasional numerically strong cohorts are able to rapidly increase declining population size.

Although this and other work (Vincenzi et al. 2012a, 2014a) have shown that marble trout populations are highly resilient to extreme events (i.e., are able to recover from major disturbances without persistent changes in structure and numbers, Gunderson 2000), complete extirpation of populations has recently occurred and represents a serious threat to the persistence of the species. Moreover, extreme rainfall and flood events are predicted to increase as a consequence of global climate change (IPCC 2007, 2012), thus further increasing the risk of extinction for marble trout. Slow-growing marble trout populations may be at greater risk of loss than fast-growing ones, as rapid growth is likely to accelerate sexual maturation and more rapidly increase fish numbers at a critical time. The creation of new populations in streams that are not affected by flash floods and debris flow is be the most effective conservation measure to increase the survival chances of the species.

Marble trout populations are largely recruitment-driven and further investigations should focus on determining the causes of variation in recruitment and movement of young. While the movement of older marble trout is limited, we do not know the extent of movement of young-of-year, which may be particularly important for the re-colonization of the stream stretches most affected by extreme events (Weese et al. 2011). Pedigree reconstruction using molecular markers will allow us to better understand the determinants of recruitment in marble trout and to quantify movement at early life stages. Finally, an important question is whether potential natural selection for faster life histories, as expected under conditions of high adult mortality or unpredictable adult environment (Stearns 1992), may increase resilience of marble trout to extreme events (Vincenzi et al. 2014a), particularly given the predicted increase in the frequency of floods.

Data, Online Code, and Supplementary Results (Online Code and Results):

http://dx.doi.org/10.6084/m9.figshare.1478066

## Acknowledgements

Simone Vincenzi is supported by an IOF Marie Curie Fellowship FP7-PEOPLE-2011-IOF for the project “RAPIDEVO” on rapid evolutionary responses to climate change in natural populations and by the Center for Stock Assessment Research (CSTAR), a partnership between University of California Santa Cruz and the Southwest Fisheries Science Center. We thank the employees and members of the Tolmin Angling Association (Slovenia) for carrying out fieldwork since 1993. We thank Travis Apgar for helping us produce Fig. 1.

